# Temporal Mapping of Radiation-Induced Neural Injury and Mitigation in Human Cortical Organoids

**DOI:** 10.64898/2026.03.04.709672

**Authors:** Ling He, Harley I. Kornblum, Aparna Bhaduri, Frank Pajonk

## Abstract

**Background:** Radiation therapy is a standard-of-care oncological treatment for central nervous system (CNS) malignancies. However, as survival outcomes improve, radiation-induced injury to normal brain tissue has increased in clinical significance. CNS radiation injury is a delayed, multifactorial process characterized by impaired neurogenesis, reactive gliosis, and persistent functional deficits. Mechanistic exploration and development of effective radiation mitigators have been limited by the lack of scalable, human-relevant models.

**Methods:** Mature human iPSC-derived cortical organoids were exposed to single-dose or clinically relevant fractionated radiation (5 x 2 Gy). DNA damage, apoptosis, and growth dynamics were assessed longitudinally. Structural organization, synaptic integrity, and neuroinflammatory responses were evaluated by immunofluorescence and real-time PCR. Transcriptomic profiling was performed at 72 hours and 2 weeks after fractionated radiation to capture acute and delayed effects. Two candidate radiation mitigators, NSPP and amisulpride, were tested for their therapeutic effects within the organoid system.

**Results:** Cortical organoids exhibited partial recovery following single doses up to 4 Gy or fractioned irradiation. Transcriptomic analyses revealed that radiation not only reduced overall cell viability but also reshaped lineage trajectories, characterized by depletion of neural stem/progenitor populations, loss of neuronal identity, enhanced gliogenesis, increased inflammatory cytokines, and disrupted cortical layering and synaptic integrity. Treatment with NSPP or amisulpride attenuated injury-associated transcriptional and structural alterations.

**Conclusion:** Human cortical organoids recapitulate key features of radiation-induced neural injury, recovery, and therapeutic modulation, providing a robust, scalable, and human-relevant platform for studying CNS radiation biology and preclinical screening of candidate radiation mitigators.

**Key points:**

- Human iPSC-derived cortical organoids enable study of human CNS radiation responses.
- Organoids recover after single-dose and fractionated radiation relevant to clinical exposure.
- The platform supports scalable, human-relevant testing of radiation mitigation strategies.

## Introduction

The terror attacks of September 11^th^, 2001, the possibility of dirty bomb attacks in the future, rising international tensions among nuclear weapon holders, renewed interest in nuclear power reactors, and the possibility of exposure to space radiation during deep space missions to the moon and beyond [1] have revived the search for effective radiation mitigators. While early efforts primarily focused on preventing acute radiation syndromes affecting rapidly dividing tissues, such as the hematopoietic and gastrointestinal systems [2, 3], there is now growing recognition of the need to address radiation-induced injury in late-responding tissues, particularly the central nervous system (CNS) [4, 5]. Likewise, established anti-cancer therapies, including chemo- and radio-therapy, impair cognitive function in women undergoing chemotherapy for breast cancer [6–8] and in pediatric patients after cranial irradiation causing a loss of 1-2 IQ points per year in the latter [9, 10]. Mitigating strategies to counteract such adverse effects arising from cancer treatments are an unmet need.

For obvious ethical and practical reasons, most studies of radiation-induced CNS injury have relied on murine models [11–15], with selected findings validated in non-human primates [16–19]. Although informative, these systems do not fully recapitulate key aspects of human brain development, cellular composition, and transcriptional regulation, limiting their translational relevance [20].

Recent advances in the field of regenerative medicine and next generation sequencing technologies have enabled the generation of increasingly complex human cortical organoids derived from induced pluripotent stem cells (hiPSCs). These three-dimensional models capture essential features of human cortical development and allow for the investigation of cellular and molecular processes at a level of resolution granularity previously impossible to achieve [21–23]. However, the utility of human brain organoids for radiobiology research is largely untested to date.

In this study, we investigated radiation responses in mature human iPSC-derived cortical organoids and evaluated their suitability as a platform for testing radiation mitigators. We demonstrated that cortical organoids recovered following single doses up to 4 Gy or clinically relevant fractioned irradiation (5 x 2 Gy), and that radiation-induced responses could be modulated by radiation mitigators previously identified in cell-based studies. Together, our findings establish human cortical organoids as a powerful and scalable model for studying CNS radiation biology and preclinical high-throughput screening platform for radiation mitigation strategies.

## Materials and Methods

### Stem Cell Culture and Maintenance

hiPSCs were maintained following established protocols consistent with previous studies [24–26]. Multiple hiPSC lines obtained from the UCLA Stem Cell Core Facility were used throughout this study **(Suppl. Table 1)**. Cells were cultured on Matrigel-coated plates in mTeSR Plus medium (Stem Cell Technologies, Vancouver, Canada) with regular medium change and passaged upon reaching approximately 80% confluency using ReLeSR (Stem Cell Technologies). Cells were routinely screened for mycoplasma contamination. Detailed culture, passaging, and cryopreservation procedures are provided in **Supplementary Materials and Methods**.

### Ethics statement

All experiments involving human iPSCs were conducted in accordance with institutional and national ethical guidelines. The use of human iPSC lines in this study was reviewed and approved by the UCLA Institutional Biosafety Committee (IBC) and Human Pluripotent Stem Cell Research Oversight Committee (hPSCRO) (UCLA IBC approval number ARC-2015-065; hPSCRP approval number 2024-001-02). All procedures complied with established protocols for the ethical handling, differentiation, and use of human pluripotent stem cells in research.

### Cortical Organoid Generation

Cortical organoids were generated using a modified protocol based on the previous publication [27], with adaptations from more recent studies [24, 26, 28]. Briefly, dissociated hiPSCs were aggregated in low-attachment plates to form embryoid bodies and cultured in stage-specific media to promote neural induction and cortical differentiation. Organoids were maintained with sequential medium formulations to support early neural patterning, neurogenesis, and later neuronal maturation. Medium changes were performed every other day, and organoids were transferred to low-attachment plates for long-term culture. Detailed media composition, small molecule treatments, and differentiation conditions are provided in **Supplementary Materials and Methods**.

### Immunofluorescence staining

Human cortical organoids were fixed in 4% paraformaldehyde, cryoprotected in sucrose, embedded in OCT, and cryosectioned (10-16 µm). Sections were subjected to antigen retrieval, permeabilization, and blocking prior to incubation with primary antibodies overnight at 4°C, followed by appropriate fluorescent secondary antibodies and nuclear counterstaining. Slides were mounted with antifade medium and imaged using a Nikon A1 confocal microscope (Nikon A1, Melville, NY). Detailed procedures, including antibody information and staining conditions, are provided in **Supplementary Materials and Methods**.

### Irradiation

Organoids were irradiated at room temperature using an experimental X-ray irradiator (Gulmay Medical Inc. Atlanta, GA) at a dose rate of 5.519 Gy/min. Control samples were sham-irradiated. The X-ray beam was operated at 300 kV and hardened using a 4 mm Be, a 3 mm Al, and a 1.5 mm Cu filter and calibrated using NIST-traceable dosimetry.

### Drug treatment

Organoids were treated with 1-[(4-Nitrophenyl)sulfonyl]-4-phenylpiperazine (NSPP, Vitascreen, Champaign, IL), or Amisulpride (#A2729, Sigma). Each compound was administered at 10 µM following 5-day-on/2-day-off schedule, beginning 24 hours after exposure to a single 8 Gy dose of radiation, and continued for up to 4 weeks.

### Western blotting

Cortical organoids were collected, washed with PBS, and lysed in ice-cold RIPA buffer (10 mM Tris-HCl (pH 8.0), 1 mM EDTA, 1% Triton X-100, 0.1% Sodium Deoxycholate, 0.1% SDS, 140 mM NaCl, 1 mM PMSF) supplemented with proteinase and phosphatase inhibitors (Thermo Fisher Scientific). Organoids were homogenized using a pellet pestle (DWK Life Science, Millville, NJ). Protein concentrations were quantified by the BCA protein assay kit (Thermo Fisher Scientific). Samples were denatured in 4x Laemmli sample buffer (Bio-Rad) containing 10% β-mercaptoethanol at 95°C for 10 minutes. Details of SDS-PAGE conditions and antibodies are provided in **Supplementary Materials and Methods**.

### Quantitative Real-time PCR

Total RNA was isolated using TRIZOL Reagent (Invitrogen), and cDNA synthesized using SuperScript Reverse Transcription IV (Invitrogen). Quantitative PCR was performed in a QuantStudio^TM^ 3 Real-Time PCR System using SYBR Green Master Mix (Applied Biosystems, Carlsbad, CA). Gene expression levels were normalized to the housekeeping gene PPIA and calculated using the ΔΔCt method. Primer sequences are listed in **Suppl. Table 2**.

### Single Cell RNA Sequencing

Day 120 cortical organoids were exposed to fractionated ionizing radiation, receiving 2 Gy per day for five consecutive days. Control organoids were sham irradiated. Organoids were harvested at two time points: 72 hours and 2 weeks following the final radiation dose. For single cell dissociation, cortical organoids were enzymatically digested into single-cell suspension via 30-min incubation at 37°C in papain and DNAse in EBSS (#LK003150, VWR), shaking vigorously every 5-10 min. Samples were then triturated with manual pipetting 10 times and pelleted with a 5-min spin at 300 *x g* in 4°C. Dissociated cells were then counted and resuspend in the provided Cell Resuspension Buffer (Fluent Bioscience, Watertown, WA), with a target capture of 50,000 cells per sample. Single cell transcriptomic profiling was performed using the PIPseq^TM^ T2 3’ Single Cell RNA Kit v5 (Fluent Bioscience).

Sequencing libraries were prepared according to PIPseq T2 instructions, with 12 cycles for cDNA amplification and 8 cycles for library amplification, depending on input DNA concentration. Final libraries were quantified using a Qubit 4 Fluorometer and TapeStation 4200, and sequenced using the NovaSeq X Plus platform following Fluent Biosciences’ specifications.

### Single Cell RNA Sequencing Data Analysis

Raw FASTQ files were aligned to the human reference genome (GRCh38) and processed using the DRAGEN Single Cell RNA app on the Illumina BaseSpace platform. Gene expression matrices were imported and analyzed using the Seurat (v5) R package [29] in RStudio. Cells expressing fewer than 500 genes or exhibiting high mitochondrial content (>5%) were excluded from further analysis. Downstream analyses, including normalization, clustering, dimensionality reduction, and cell-type annotation, were conducted following established pipelines as described in [30]. Trajectory analyses were performed using the R package monocle3 (version 1.4.26) [31]. Senescence scores were calculated using the Python package SenePy [32].

### Statistics

All data shown are represented as mean ± standard error mean (SEM) of at least 3 biologically independent experiments. A *p*-value of ≤0.05 in an unpaired two-sided *t*-test or one-sided ANOVA followed by Tukey’s multiple comparison test indicated a statistically significant difference.

## Results

### Establishment and Characterization of Functionally Mature Human Cortical Organoids

To establish a physiologically relevant platform that recapitulates human brain structures and microenvironment, we generated cortical organoids from hiPSCs using an established protocol [27] **(Fig. 1A)**. Human iPSCs were maintained as undifferentiated colonies in mTeSR medium **(Fig. 1B)**. After transfer to V-bottom 96-well plates and treatment with ROCK, TGF-β, and Wnt inhibitors, embryoid bodies (EBs) with forebrain identity were formed. By day 18, uniform spheroid aggregates were consistently observed **(Fig. 1C)**, and these continued to expand in size and structural complexity over time. By week 5, rosette-like structures characteristic of early cortical development were evident in brightfield microcopy **(Fig. 1D)**. In total, we have tested 10 patient-derived iPSC lines (7 female, 3 male), with only 5 lines (4 female, 1 male – highlighted bold in **Fig. 1E**) reliably generating cortical organoids of consistent size (∼2-3 mm), which were used for subsequent experiments.

**Figure 1.**
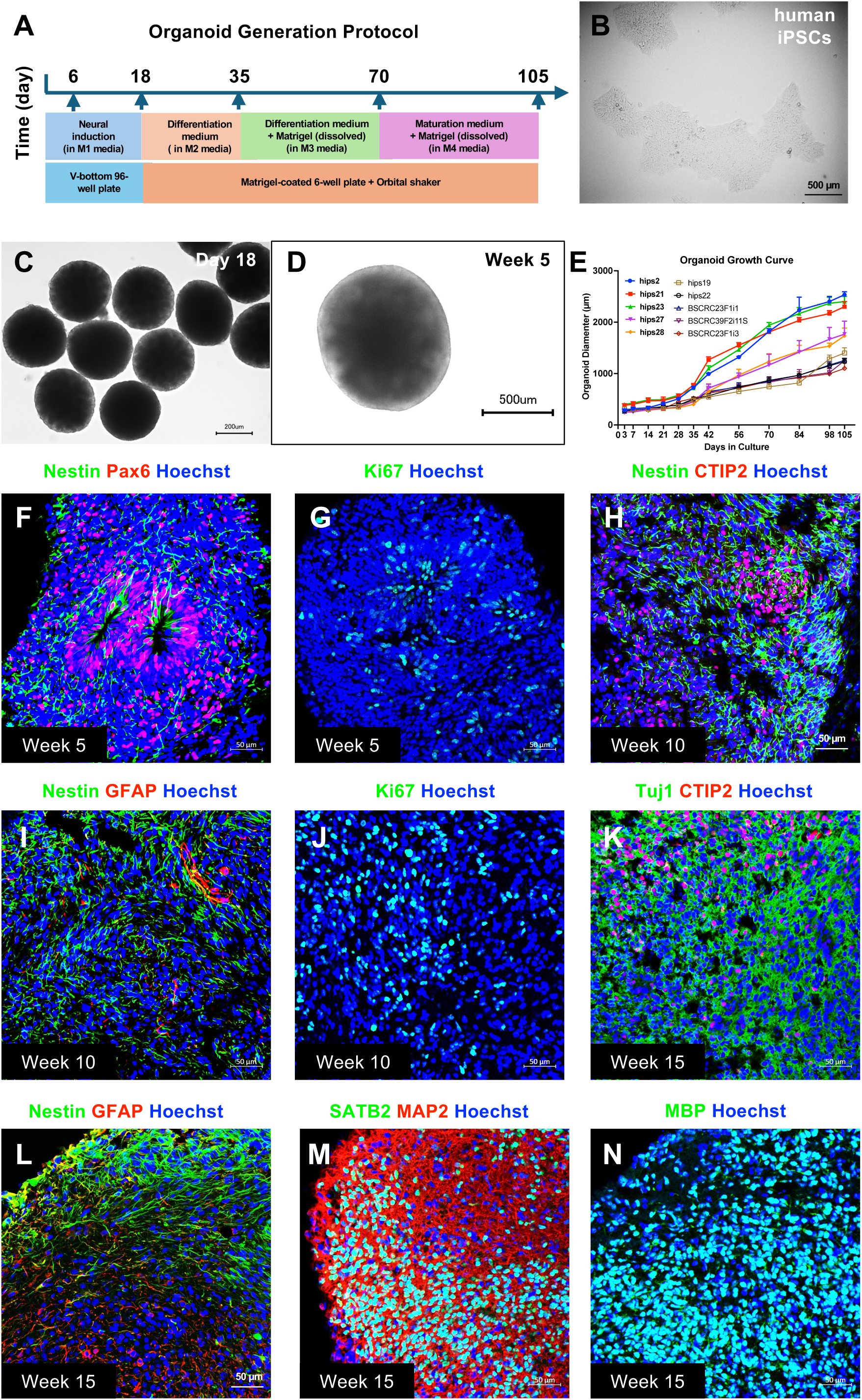
Establishment and characterization of functionally mature human iPSC-derived cortical organoids. **(A)** Schematic timeline of the organoid generation protocol from human iPSCs, including neural induction (day 0-6), differentiation (day 18-35), and maturation stages (day 70 onward). **(B)** Representative phase-contrast image of human iPSCs prior to neural induction. **(C/D)** Brightfield images showing organoid morphology at day 18 **(C)** and week 5 **(D)**, with consistent spherical growth and rosette structure formation. **(E)** Growth curve of organoids derived from multiple human iPSC lines, demonstrating reproducible increase in diameter over time (the bold highlighted lines were included in the subsequent experiments, while the remaining ones showed only modest increase of their organoid sizes during culture). **(F/G)** Immunofluorescence staining confirms neural lineage development by Nestin⁺/Pax6⁺ neural progenitors staining and Ki67⁺ proliferating cells stained at week 5. **(H-J)** Immunofluorescence staining of neural stem/progenitor cell marker Nestin, deep-layer neuron marker CTIP2, astroglial cell marker GFAP, and Ki67 at week 10. **(K-N)** Immunofluorescence staining of Nestin, GFAP, CTIP2, upper-layer excitatory neuron marker SATB2, mature neuron marker MAP2, and mature oligodendrocyte marker MBP at week 15.

To define cellular composition of cortical organoids during development, we performed immunofluorescence staining at sequential stages. At week 5, organoids displayed rosette-like structures enriched in Nestin+ and Pax6+ neural stem/progenitor cells, along with abundant Ki67+ proliferative cells **(Fig. 1F/G)**. By week 10, the stem/progenitor pool began to differentiate, giving rise to CTIP2+ deep-layer cortical neurons, while GFAP+ astroglial populations emerged, indicating that neurogenesis and gliogenesis occur concurrently **(Fig. 1H/I)**. Proliferation persisted at this stage, as evidenced by Ki67+ cells **(Fig. 1J)**. By week 15, organoids exhibited greater neuronal complexity, including Tuj1+ and CTIP2+ neurons, SATB2+ upper-layer neurons, and MAP2+ mature neuronal populations which integrated into organized structures **(Fig. 1 K-M)**. Importantly, MBP+ oligodendrocytes were also detected at this stage **(Fig. 1N)**, demonstrating myelination capacity, a key hallmark of functional maturation. Taken together, these findings show that human iPSC-derived cortical organoids recapitulate key features of *in vivo* cortical development through maturation and validate this platform as a powerful model for evaluating radiation-induced injury in a human-relevant setting.

### Radiation-Induced DNA Damage, Apoptosis, and Alterations in Organoid Growth Dynamics

To assess the impact of radiation on organoid development, we irradiated mature cortical organoids at day 120 and monitored organoid size over time following exposure to a range of radiation doses (0, 2, 4, 6, 8 Gy, or fractionated 5 x 2 Gy, reflecting clinically relevant fractionated regimens). Day 120 organoids were selected to better represent advanced neuronal maturation and glial development, thereby allowing evaluation of radiation effects in a more developmentally mature neural context. Growth curve analysis demonstrated a dose-dependent reduction in organoid expansion across both male- and female-derived lines **(Fig. 2A)**. Higher doses (6 Gy and 8 Gy) exhibited the most pronounced reduction in organoid size, while fractionated radiation also impaired growth but to a lesser extent compared with a single dose of 6 Gy or 8 Gy. Notably, based on the α/β ratio for brain tissue, 5 x 2 Gy fractionated radiation is biologically equivalent to ∼6 Gy delivered as a single dose.

**Figure 2.**
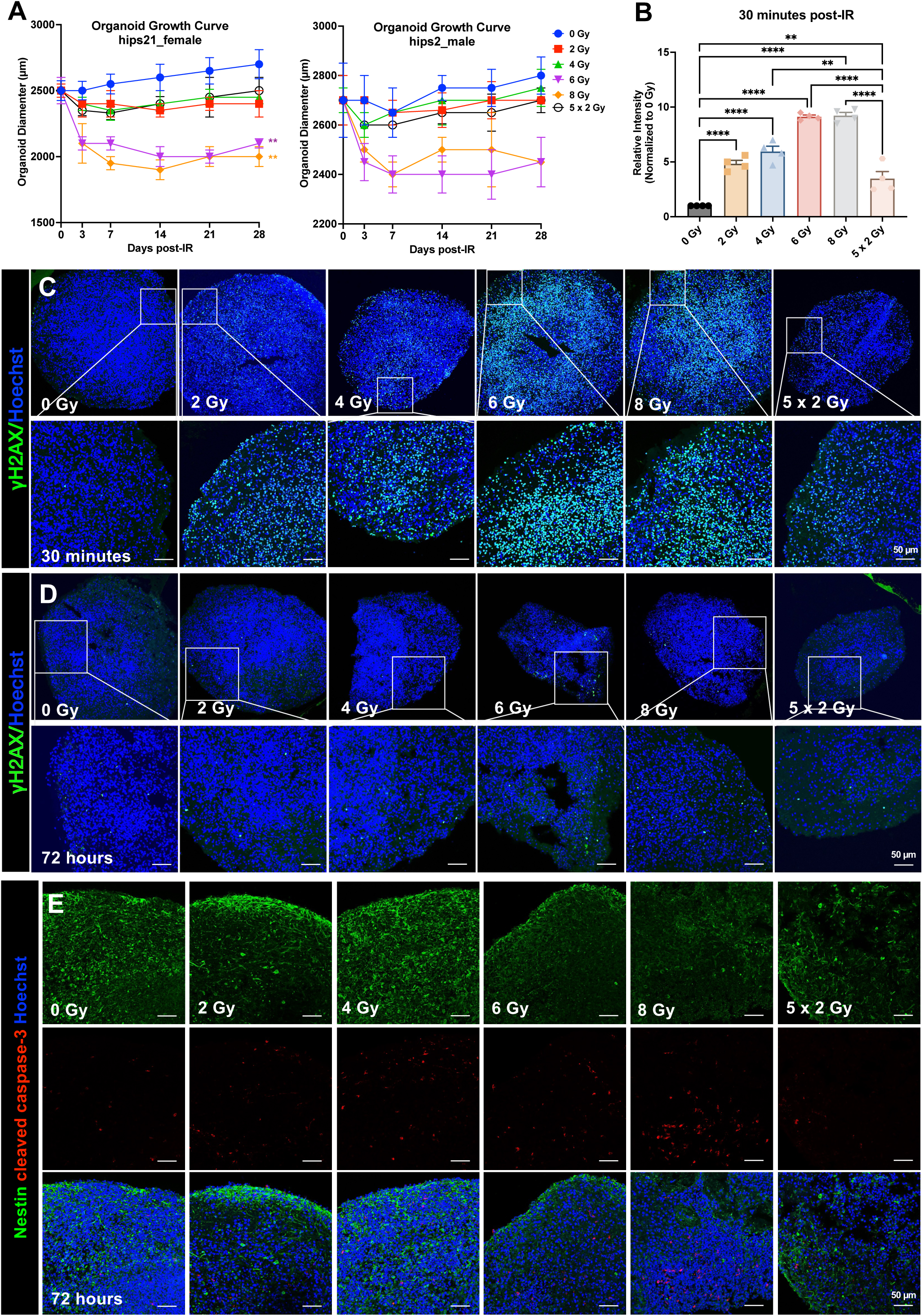
The effects of radiation on DNA damage, apoptosis, and organoid growth. **(A)** Growth curves of cortical organoids derived from both female and male iPSC lines following exposure to increasing doses of radiation (0, 2, 4, 6, 8 Gy, or fractionated 5 × 2 Gy). **(B/C)** Representative γH2AX immunofluorescence images at 30 minutes post-irradiation demonstrating dose-dependent induction of DNA double-strand breaks, with quantification performed using ImageJ. **(D)** γH2AX staining at 72 hours post-irradiation shows efficient DNA repair across all radiation conditions. **(E)** Representative immunofluorescence staining of Nestin and cleaved caspase-3 at 72 hours post-irradiation illustrating persistent apoptotic activity and associated damage to Nestin-positive neural stem/progenitor cell populations. All experiments have been performed with at least 3 biological independent repeats. *P*-values were calculated using one-way ANOVA for B. ** *p*-value < 0.01, *** *p*-value < 0.001, **** *p*-value < 0.0001.

DNA double-strand breaks are a well-known consequence of radiation exposure, but it remained unclear how this response would be reflected in our cortical organoid system and to what extent different radiation doses would induce such damage. To address this, we assessed γH2AX expression levels at early (30 minutes) and delayed (72 hours) time points post-irradiation. At 30 minutes, organoids displayed a clear dose-dependent increase in γH2AX signal intensity, with the extent of γH2AX foci being highest following 6 Gy and 8 Gy exposure, with fractionated radiation producing an intermediate signal when compared to a single dose of 6 Gy or 8 Gy **(Fig. 2B/C)**. By 72 hours post-irradiation, γH2AX levels had markedly decreased across all groups, indicating efficient DNA repair **(Fig. 2D)**. Though some residual foci persisted in higher-dose conditions (6 Gy and 8 Gy), no significant differences were observed between doses. Collectively, these results demonstrate that radiation induces rapid and dose-dependent DNA damage in cortical organoids, but that efficient DNA repair mechanisms resolve most damages by 72 hours.

To further investigate the mechanisms underlying the suppression of organoid growth dynamics, we next examined the effects of radiation on apoptosis. At 72 hours post-irradiation, we observed a dose-dependent increase in cleaved caspase-3 positive cells, indicating enhanced apoptotic activity with higher radiation doses. Notably, fractionated irradiation resulted in substantially fewer apoptotic cells compared to the biologically equivalent single dose (∼6 Gy), suggesting reduced acute cytotoxicity under fractionated conditions **(Fig. 2E)**. We also co-stained for Nestin, a marker of neural stem/progenitor cells, as they have been known for their sensitivity/vulnerability to radiation. Radiation exposure did lead to a marked reduction in Nestin+ cells, however, cleaved caspase-3 positive cells did not co-localize with the Nestin+ population. This pattern suggests that classical apoptosis alone may not fully account for the reduction of Nestin+ population. It remains possible that a subset of progenitor cells underwent rapid cell death prior to detection, or alternatively, that radiation impaired their self-renewal capacity and maintenance of stem-like identity.

### Temporal Dynamics of Radiation-Induced Transcriptional Changes in Cortical Organoids

To investigate whether radiation impacts lineage specification and function in our cortical organoid system, we performed real-time PCR to profile gene expression dynamics across multiple radiation doses. We included 72 hours to capture acute responses, and 2 and 4 weeks to assess delayed effects. A heatmap of representative lineage-associated and cortical layer-specific genes is shown in **Fig. 3A**. We observed a consistent, dose-dependent reduction in neural stem/progenitor cell markers, including Nestin, Sox2, and Pax6, as early as 72 hours post-radiation, consistent with the well-established radiosensitivity of proliferative and self-renewing populations. This reduction persisted through the 2-week time point, indicating a sustained disruption of neural lineage maintenance. By 4 weeks, expression of these markers began to recover, suggesting that residual or newly emerging progenitor-like cells may partially replenish the progenitor pool over time. In parallel, neuronal lineage genes such as TUBB3 (βIII-tubulin) and MAP2 showed mild suppression across radiation groups, suggesting that radiation not only depletes neural progenitors but also impairs neuronal differentiation.

**Figure 3.**
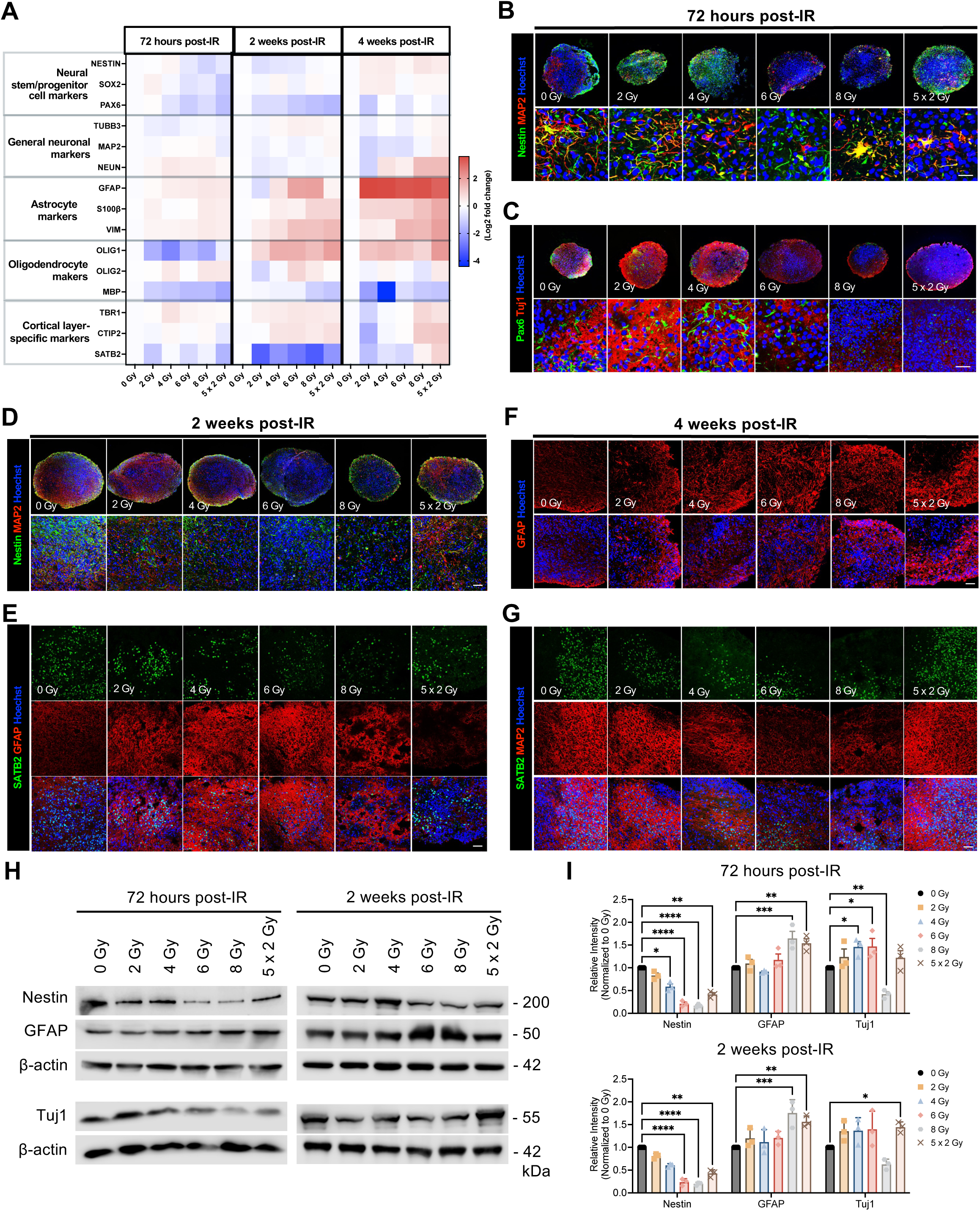
Radiation-induced transcriptional changes and disruption of structural organization in cortical organoids. **(A)** Heatmap showing representative gene expression profiles across different radiation doses and time points, measured by real-time PCR. **(B/C)** Immunofluorescence staining of Nestin and MAP2, and of Pax6 and Tuj1, at 72 hours following exposure to increasing doses of radiation (0, 2, 4, 6, 8 Gy, or fractionated 5 × 2 Gy). **(D/E)** Immunofluorescence staining of Nestin and MAP2, and of SATB2 and GFAP, at 2 weeks post-irradiation under the same dose conditions. **(F/G)** Immunofluorescence staining of GFAP, and of SATB2 and MAP2, at 4 weeks after radiation exposure. Scale bar: 50 µm. **(H/I)** Western blot analysis of Nestin, GFAP, Tuj1 at 72 hours and 2 weeks post-irradiation, with β-actin serving as the loading control. Quantification was performed using ImageJ. All experiments have been performed with at least 3 biological independent repeats. *P*-values were calculated using one-way ANOVA for I. * *p*-value < 0.05, ** *p*-value < 0.01, *** *p*-value < 0.001, **** *p*-value < 0.0001.

Intriguingly, we found that glial markers, such as GFAP, S100β, vimentin, Olig1, and Olig2 were markedly upregulated, particularly in organoids exposed to higher radiation doses, indicating a shift toward gliogenesis and the emergence of reactive glial phenotypes. This effect was most pronounced at 2 and 4 weeks post-radiation, indicating a sustained and progressive remodeling of lineage fate over time. Meanwhile, expression of TBR1 and CTIP2, which mark deep-layer cortical neurons, was increased following irradiation, whereas expression of SATB2, a marker of upper-layer neurons, was consistently reduced. This shift implies that radiation disrupts cortical layer organization in the organoids. Because deep-layer neurons are generated earlier and upper-layer neurons later during corticogenesis, the observed changes likely depend on the developmental stage at which irradiation occurs and may reflect altered maintenance or maturation of these neuronal populations, which still may have important implications for cortical circuit organization and long-range connectivity. Together, these transcriptional profiles demonstrate that radiation not only reduces overall cell viability but also actively reshapes lineage trajectories in cortical organoids. The observed pattern, loss of stem/progenitors and neuronal identity, along with enhanced gliogenesis and disruption of cortical layering, suggests a coordinated response that may explain the long-term neurodevelopmental deficits and cognitive impairments seen following radiation exposure.

### Impact of Radiation on Structural Organization, Synaptic Integrity, and Neuroinflammatory Responses in Cortical Organoids

To further evaluate how radiation affects the structural organization of mature cortical organoids, we performed immunofluorescent staining using lineage-specific markers, including Nestin and Pax6 (neural stem/progenitors), MAP2 and Tuj1 (neuronal differentiation and maturation), GFAP (astrocytes), and SATB2 (upper-layer cortical neurons). Consistent with the transcriptional changes, we observed a reduction in progenitor marker expression, as well as neuronal differentiation and maturation marker expression **(Fig 3. B-E)**. In contrast, GFAP staining was strongly enhanced, while SATB2 expression was notably decreased **(Fig. 3E-G)**, further suggesting the shift to gliogenesis and potential disruption of cortical layering after radiation.

Synaptic integrity and neural connectivity are obviously disrupted by radiation, a primary cause of radiation-induced cognitive decline in cancer patients receiving cranial radiotherapy [33, 34]. To evaluate if our model could recapitulate those effects upon radiation exposure, we examined the expression of some key pre- and post-synaptic markers in cortical organoids. Real-time PCR analysis revealed a modest decrease in SYN1 (Synapsin 1) and PSD1 (postsynaptic density protein 1) expression across increasing radiation doses, indicating impaired synaptic maturation and reduced synaptic density **(Fig. 4A)**. However, these decreases were considerably less pronounced in the fractionated irradiation group, particularly for SYN1, suggesting that fractionated dosing may partially preserve synaptic integrity compared with an equivalent single-dose regimen. Further, these transcriptional changes were confirmed by immunofluorescence staining, which demonstrated a marked reduction in Synapsin 1 at both acute (72 hours) and chronic (4 weeks) time points **(Fig. 4B/C)**. In contrast, organoids exposed to 5 x 2 Gy retained more Synapsin-1+ structures, supporting a potential protective effect of fractionated radiation on synaptic maintenance.

**Figure 4.**
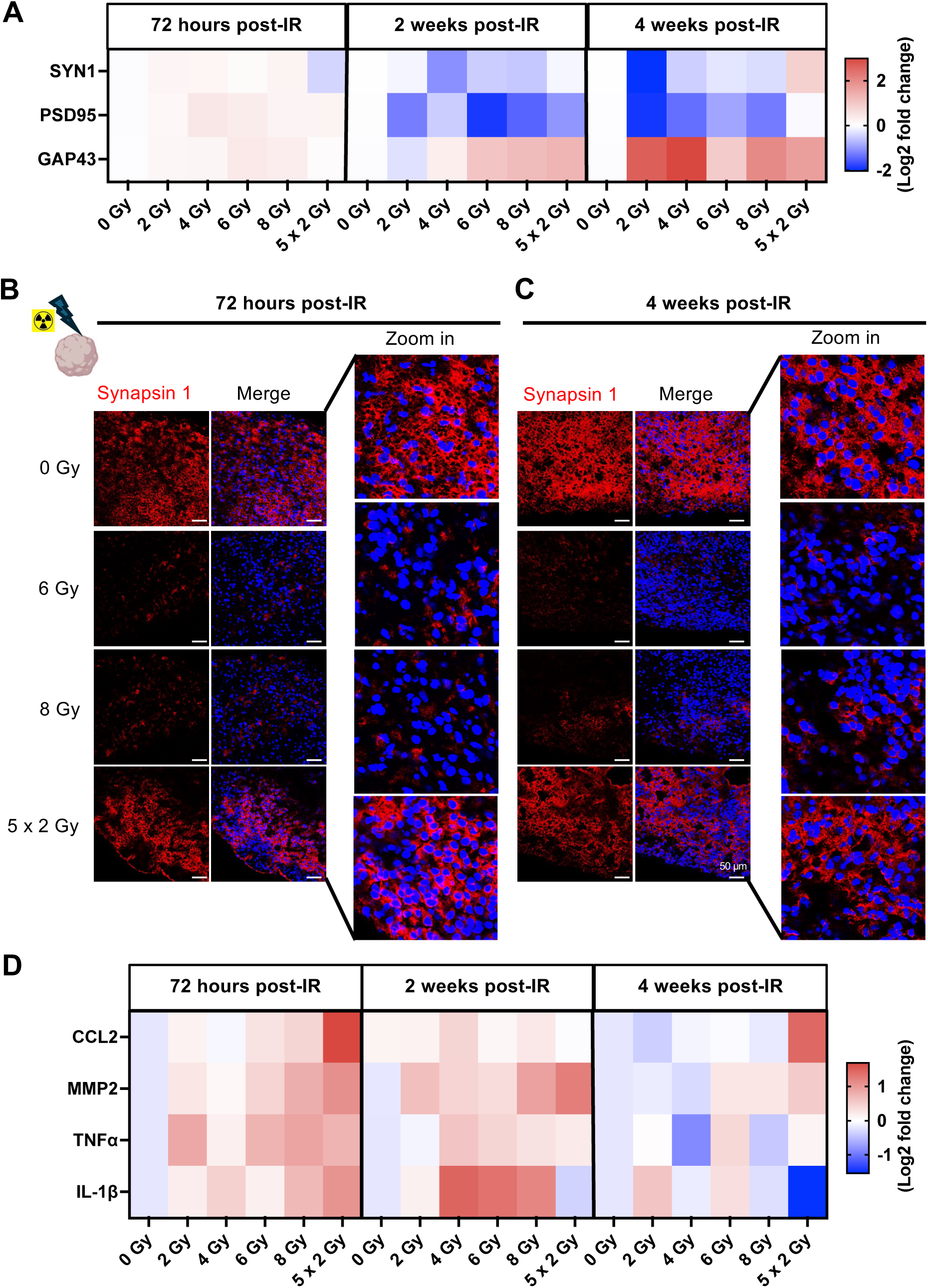
Radiation-induced synaptic impairment and neuroinflammation in cortical organoids. **(A)** Heatmap showing synaptic gene expression profiles across radiation doses and time points, measured by real-time PCR. **(B/C)** Immunofluorescence staining of Synapsin 1 at 72 hours and 4 weeks following exposure to radiation (0, 6, 8 Gy, or fractionated 5 × 2 Gy). **(D)** Heatmap showing representative neuroinflammatory gene expression profiles across radiation doses and time points, measured by real-time PCR. All experiments have been performed with at least 3 biological independent repeats.

Intriguingly, radiation induced a robust, dose-dependent increase in GAP43, which is a marker of neurite outgrowth and axonal remodeling **(Fig. 4A)**. Its upregulation may reflect a compensatory regenerative attempt by neurons to re-establish structural connectivity following radiation-induced synaptic disruption.

Lastly, to assess whether these synaptic alterations were accompanied by neuroinflammatory activation, we profiled the expression of pro-inflammatory cytokines and chemokines across radiation doses and time points. Transcriptional analysis showed robust, dose-dependent increases in CCL2, TNF-α, IL-1β, and the extracellular matrix-modifying enzyme MMP2 **(Fig. 4D)**. These changes are consistent with the establishment of a pro-inflammatory microenvironment known to exacerbate neuronal stress, impair synaptic recovery, and potentially contribute to the long-term cognitive decline.

### Fractionated Radiation Induces Distinct Cell-Type-Specific Transcriptomic Changes as Evidence by Single-Cell RNA Sequencing

Given the protective effects of fractionated radiation observed in our cortical organoid model, as well as its clinical relevance, we next performed single-cell RNA sequencing (scRNA-seq) on mature cortical organoids exposed to 5 x 2 Gy fractionated regimen to study the radiation effects on cell fate in more detail. We included two time points, 72 hours post-irradiation representing the acute phase, while 2 weeks post-irradiation representing the chronic phase. The 2-week time point was selected because our earlier experiments revealed the most pronounced and distinct structural and molecular alterations at this stage compared to the later, 4-week post-irradiation time point.

The UMAP visualization by sample **(Fig. 5A)** showed that the overall cellular landscape largely overlapped at the transcriptional level across treatment groups. Annotation confidence was high across most clusters **(Fig. 5B)**, indicating that radiation did not generate entirely new cell populations but instead altered the relative abundance of existing ones. Lovain clustering identified 10 unique cell clusters that were represented across the different treatment arms of the study **(Fig. 5C)**. We next compared marker genes of the identified clusters to established gene expression signatures from the adult and developing brain, as previously described [7]. In untreated cortical organoids, the predominant populations consisted of excitatory neuron (52.6 %), radial glia (RG; 19 %), newborn neurons (8.3 %), dividing cells (4.5 %), inhibitory neurons (3.9 %), and dividing radial glia (RG. Div; 3 %). 72 hours after the 5 x 2 Gy fractionated radiation regimen, the most notable change was a marked reduction in proliferative populations, with dividing cells decreasing to 0.28 % and RG dividing cells nearly eliminated (0.07 %). This was accompanied by an increase in the excitatory neuron population to 66.1 % and the emergence of an endothelial-like population (9 %). At 2 weeks post-radiation, proliferative cells remained suppressed (0.5 %), while excitatory neurons continued to dominate (65.2 %), and the endothelial population was reduced to 2.6 % **(Fig. 5D; Suppl. Table 3)**.

**Figure 5.**
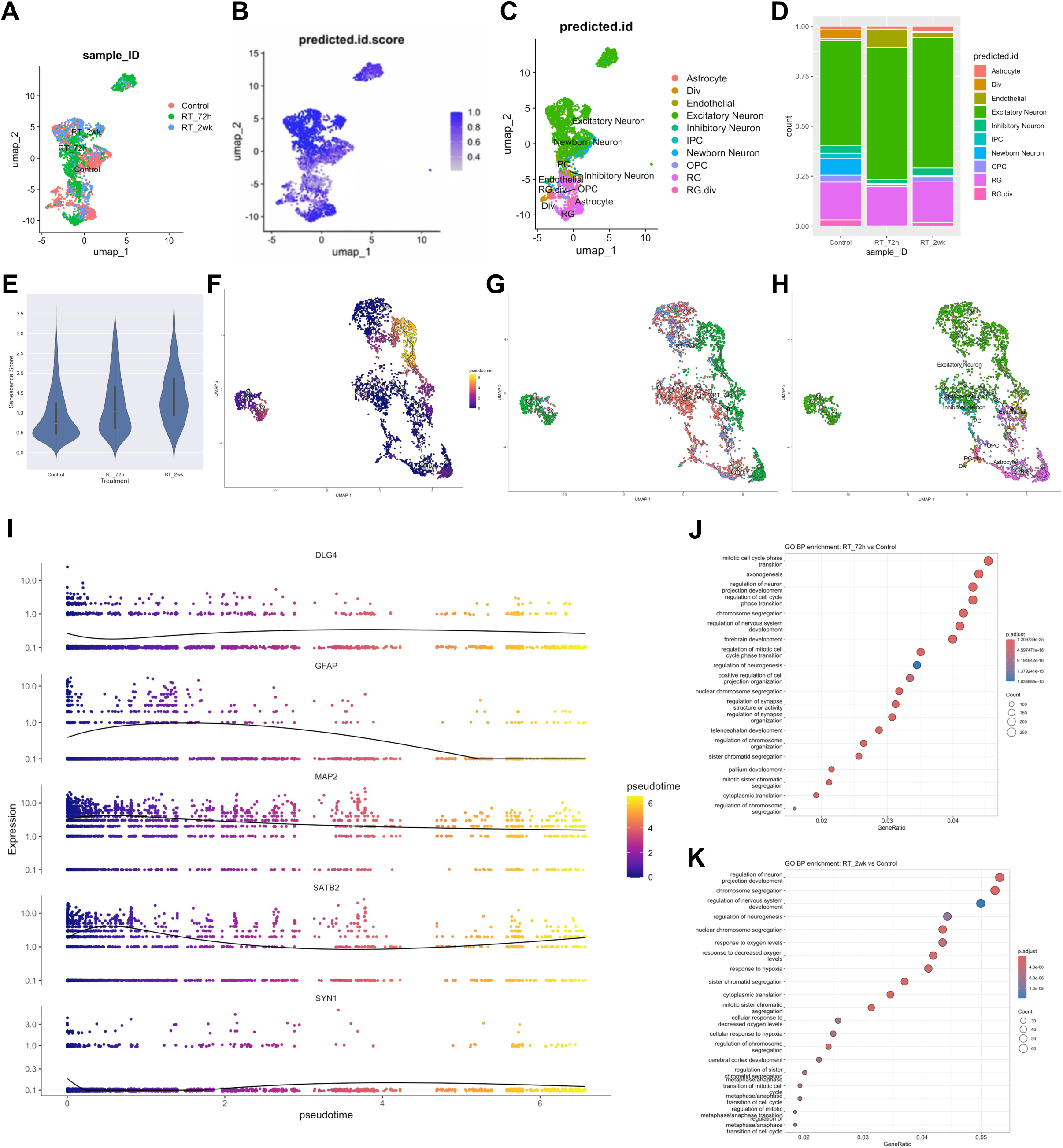
Fractionated radiation induces distinct cell-type-specific transcriptomic changes, and senescence, as determined by Single-Cell RNA Sequencing. **(A-C)** UMAP plots of clusters identified 72 hours and 2 weeks after the 5 x 2Gy fractionated radiation regimen (RT), along with the predicted annotation confidence score for clusters and projected cell type annotations. **(D)** Stacked column graph showing the relative distribution of identified cell types across all treatment groups. **(E)** Violin plot illustrating senescence scores across treatment conditions. **(F)** scVelo dynamical model latent time projected onto the UMAP embedding, illustrating inferred transcriptional state progression. **(G)** scVelo latent time colored by treatment condition, showing differential distribution of control and irradiated cells along the inferred trajectories. **(H)** scVelo latent time overlapped with cell-type annotations, demonstrating that velocity-defined state progression occurs within established neural and progenitor populations. **(I)** Expression of representative neuronal and glial markers (DLG4, GFAP, MAP2, SATB2, SYN1) plotted along pseudotime inferred from the scVelo dynamical model. **(J/K)** Gene Ontology (GO) biological process enrichment analysis of differentially expressed genes comparing RT_72h versus Control and RT_2wk versus Control.

To explore the radiation effects on cell senescence, we applied a universal cellular senescence signature for scoring [32]. Compared with control organoids, radiation exposure resulted in a clear induction of senescence, with the highest score observed at 2 weeks post-radiation, indicating a sustained induction of senescence-related states following radiation **(Fig. 5E)**. Considering the alterations in cluster composition between different treatments we used dynamical model to compute cell trajectories based on RNA expression and splicing information by calculating RNA velocities, dynamical genes, and latent time [35] for the two different treatment timepoints and the control sample cells that were used as the starting point. UMAP plots of latent time suggested that control organoid cells originated from newborn neurons and excitatory neurons, while the irradiated organoids branched from excitatory neurons and endothelial-like cells **(Fig. 5 F-H)**. We further validated several gene expression dynamics plotted along pseudotime for representative neuronal and glial markers **(Fig. 5I)**. Cells are ordered according to their inferred transcriptional progression, with color indicating increasing pseudotime. Neuronal markers, including DLG4, MAP2, SATB2, and SYN1, showed gradual changes in expression along pseudotime, consistent with transitions across neuronal differentiation and maturation states captured by the velocity model. GFAP exhibits a distinct expression pattern along pseudotime, reflecting heterogeneity in glial-associated or non-neuronal transcriptional states within the inferred trajectory. We also listed out the top 20 DEGs plotted along pseudotime inferred from the scVelo dynamic model **(Suppl. Figure 1)**.

Lastly, we performed Gene Ontology (GO) biological process enrichment for differentially expressed genes in irradiated samples relative to controls at 72 hours and 2 weeks post-radiation. At 72 hours, enriched GO terms are primarily associated with mitotic cell cycle regulation, chromosome segregation, and nuclear division, indicating transcriptional programs related to proliferative and cell-cycle-associated processes **(Fig. 5J)**. At 2 weeks post-radiation, enriched GO terms shift toward categories related to nervous system development, neuronal differentiation, synaptic organization, and axon-related processes, reflecting changes in gene expression associated with differentiation and maturation within neural lineages **(Fig. 5K)**.

### Assessing the Effects of Radiation Mitigators Using Human Cortical Organoids

Finally, we evaluated whether this hiPSC-derived cortical organoid platform could be utilized to assess the efficacy of radiation mitigators. A single radiation dose of 8 Gy was selected, as it has been shown to induce robust changes in neuronal structure and function. We tested two candidate mitigators, NSPP and Amisulpride, both of which we have demonstrated previously to possess mitigation activity in the gut and brain [36–38]. Mature cortical organoids were exposed to a single dose of 8 Gy, and treatment with 10 µM NSPP or 10 µM Amisulpride was initiated 24 hours post-radiation at 5-days-on/2-days-off schedule for 2 weeks **(Fig. 6A)**. Organoid growth was monitored by measuring their size, as demonstrated in **Fig. 2**. Radiation exposure resulted in a significant reduction in organoid size, whereas treatment with either NSPP or Amisulpride significantly attenuated their growth inhibition **(Fig. 6B)**.

**Figure 6.**
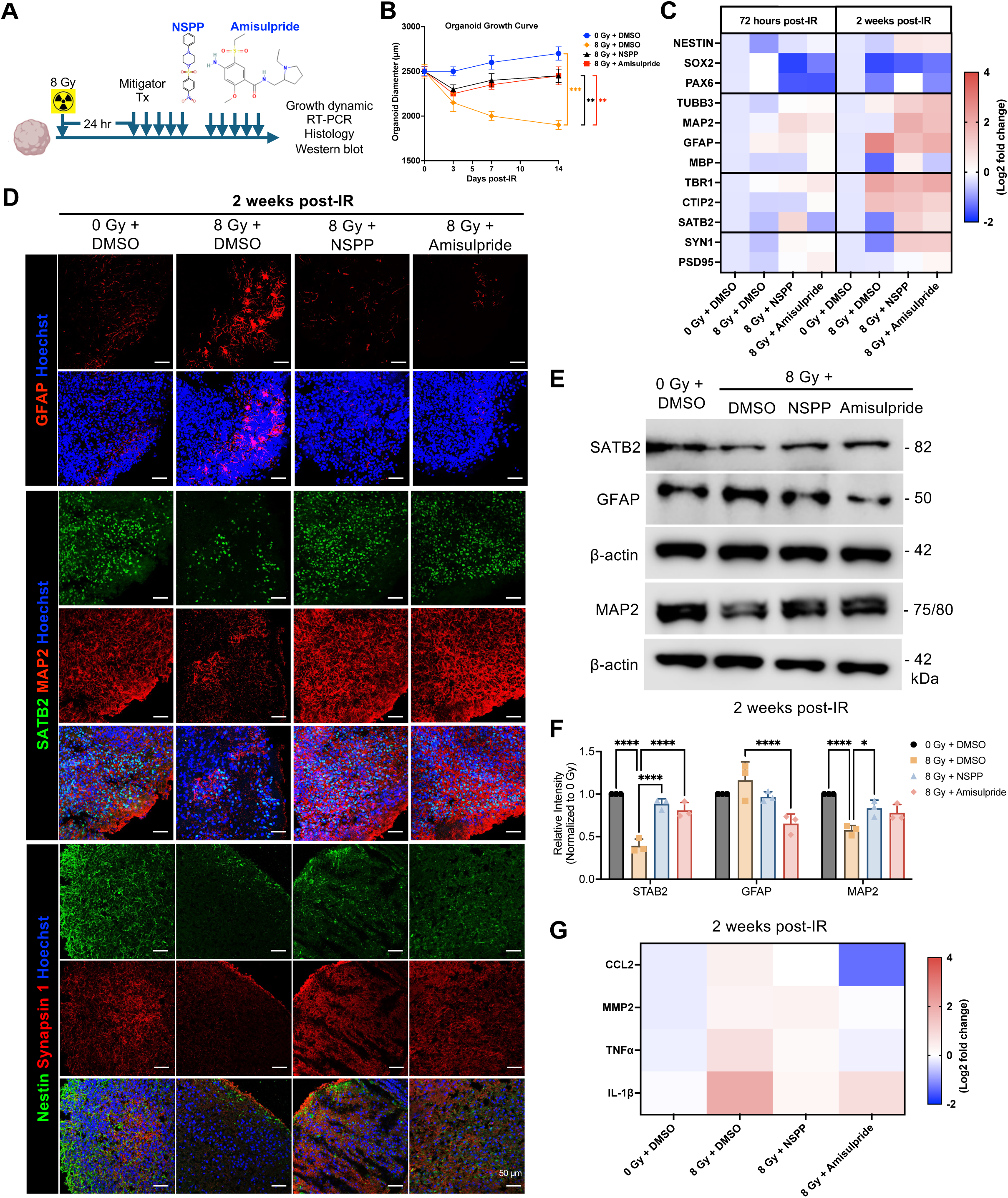
Evaluation of radiation mitigators in human cortical organoids. **(A)** Schematic overview of the experimental design for radiation mitigation studies using human cortical organoids. **(B)** Growth curve of hiPSCs-derived cortical organoids following exposure to a single dose of 8 Gy radiation, with or without treatment with 10 µM NSPP or 10 µM Amisulpride. **(C)** Heatmap showing representative gene expression profiles across mitigator treatments and time points, as measured by quantitative real-time PCR. **(D)** Immunofluorescence staining of GFAP, SATB2 and MAP2, and Nestin and Synapsin 1 in organoids at 2 weeks after a single dose of 8 Gy radiation exposure, in the presence or absence of NSPP or Amisulpride treatment. **(E/F)** Western blot analysis of SATB2, GFAP, and MAP2 at 2 weeks post-radiation with or without NSPP or Amisulpride treatment. β-actin was used as the loading control. Quantification was performed using ImageJ. **(G)** Heatmap showing representative neuroinflammatory gene expression profiles across all experimental conditions. All experiments have been performed with at least 3 biological independent repeats. *P*-values were calculated using one-way ANOVA for B. * *p*-value < 0.05, ** *p*-value < 0.01, *** *p*-value < 0.001, **** *p*-value < 0.0001.

To further assess whether these mitigators could modulate radiation-associated changes in neural stem/progenitor populations, neuronal differentiation, cortical layer specification, as well as the synapses, we performed quantitative real-time PCR analysis of representative genes at both 72 hours and 2 weeks post-radiation. Expression of NES, TUBB3, MAP2, SATB2, and SYN1 was reduced following radiation and was partially restored by both NSPP and Amisulpride at both time points, with more pronounced effects observed at 2 weeks. In addition, radiation-associated increase in GFAP expression was substantially reduced by both treatments, suggesting their attenuation of gliosis-related changes **(Fig. 6C)**. Mitigation-associated effects were further supported by immunofluorescence staining of organoids at 2-week post-radiation **(Fig. 6D)** and western blot analysis **(Fig. 6E/F)**. Finally, treatment with NSPP or Amisulpride was associated with reduced expression of radiation-induced pro-inflammatory genes, including TNFα, IL-1β, CCL2, and MMP2 **(Fig. 6G)**, indicating the potential suppression of neuroinflammation.

## Discussion

In this study, we establish human iPSC-derived cortical organoids as a robust and scalable platform for investigating CNS responses to ionizing radiation and evaluating candidate radiation mitigators in a human-relevant context. By integrating fractionated radiation regimen, single-cell transcriptomic profiling, trajectory inference, and mitigation studies, we demonstrate that mature human cortical organoids recapitulate key features of radiation-induced neural injury, recovery, and modulation that have been challenging to model using existing systems.

Historically, radiobiology research has primarily focused on acute radiation syndromes in rapidly dividing tissues, including the hematopoietic and gastrointestinal systems [39]. In contrast, radiation-induced injury to the CNS develops over time and involves sustained effects on neural progenitors, differentiation, synaptic integrity, and inflammatory signaling pathways [40–43]. Most of the current understanding of these processes is derived from murine models, which carry major drawbacks including the large number of animals required and fundamental interspecies differences in brain structures, such as variations in radial glia organization and cortical layering [20, 44] that may impact injury responses and limit direct translational relevance. Furthermore, research involving non-human primate (NHP) models is becoming increasingly difficult and costly to perform. Our findings help bridge this translational gap by demonstrating that human cortical organoids retain sufficient cellular diversity and developmental organization to capture both early DNA damage responses and delayed lineage-specific alterations following radiation within a human-derived neural context.

Compared with conventional two-dimensional human neural cell culture, cortical organoids preserve three-dimensional architecture, multicellular interactions, and most importantly the spatial organization, all of which impact radiation sensitivity, subsequent recovery, and delayed radiation effects [45, 46]. While pure two-dimensional cultures are more valuable for dissecting cell-autonomous responses [47], organoids enable analysis of radiation-induced perturbations withing coordinated neurodevelopmental trajectories and allow identification of lineage-specific vulnerabilities across progenitors, differentiating neurons, and more mature neuronal populations. In contrast, although traditional murine or NHP models provide a native brain microenvironment, the interspecies differences of brain structures such as the radial glia and inner fiber layer are undeniable [20, 48], limiting their direct translational relevance.

Recent work investigating radiation effects in human cortical organoids has largely centered on differences between photon and proton modalities, particularly in the context of pediatric brain tumors [49]. By comparing conventional photons, plateau protons, and spread-out Bragg peak (SOBP) protons both shared and distinct transcriptional alterations were identified. Notably, SOBP protons were reported to downregulate genes more involved in neurodevelopment, synaptic activity, and network function more when compared to the other methods, suggesting distinct biological consequences arise depending on radiation delivery method. In contrast, our study focuses more broadly on the biological effects of ionizing radiation exposure in human cortical organoids as a platform to model CNS injury by assessing radiation-induced changes in neural lineage trajectories, differentiation dynamics, and long-term functional maturation, and develop potential mitigation strategies. Furthermore, while earlier studies have largely relied on bulk or limited transcriptomic analyses [49], our use of single-cell transcriptomic profiling combined with trajectory inference analysis allowed us to resolve cell type-specific vulnerabilities and shifts in developmental state. These effects can be observed over time, extending beyond immediate apoptosis or cell loss.

To our knowledge, this study provides one of the first single-cell transcriptomic analyses of mature human cortical organoids in response to ionizing radiation. Consistent with prior reports of CNS radiation injury, we observed transcriptional signatures indicative of reactive gliosis following radiation, including upregulation of astrocyte-associated markers, and inflammatory cytokines. Reactive gliosis is a well-recognized feature of radiation-induced brain injury, which is an inflammatory response characterized by the proliferation of microglia and astrocytes [50–52]. The presence of these signatures in our organoid model further supports the capacity of this platform to recapitulate key aspects of human CNS injury. In addition, we observed the emergency of an endothelial-like cell population upon radiation exposure in our sequencing data, despite the absence of *bona fide* endothelial cells in standard cortical organoid culture. This finding suggests radiation-induced cellular plasticity rather than expansion of a pre-existing vascular lineage. One possibility could be stress-induced transdifferentiation or partial lineage reprogramming under genotoxic conditions, which requires further investigation.

The mitigation effects observed in our study are consistent with the reported mechanisms of action of the candidate compounds. NSPP is a Smoothened receptor agonist that we have previously characterized in both brain and GI systems [36, 37]. Pharmacologic activation of the Sonic Hedgehog (Shh) signaling pathway through Smoothened receptor agonists, such as purmorphamine and Smoothened agoinist (SAG), has been shown to exert neuroprotective and regenerative properties in injuries like ischemic stroke [53, 54], subarachnoid hemorrhage [55], and traumatic brain injury [55, 56]. Activation of Shh signaling reduces neuronal apoptosis, mitigates brain edema, and supports tissue repair. In parallel, amisulpride, a selective D2/D3 receptor antagonist, shows emerging evidence of neuroprotective and regenerative properties beyond its primary use in schizophrenia and dysthymia [57, 58]. Dopaminergic signaling impacts neural stem/progenitor dynamics, synaptic plasticity, and neuroinflammatory responses. Amisulpride has been reported to modulate microglia activation, reduce tau hyperphosphorylation [59], and enhance cognitive function [58]. Moreover, in our prior *in vivo* work, amisulpride combined with radiation significantly increased the number of neural stem/progenitor cells [38], suggesting a potential role in preserving or restoring neurogenic capacity under genotoxic stress. Together, these findings provide mechanistic support for the mitigation effects we observed in our cortical organoid system.

Despite the advantages shown here for our organoid platform, several limitations should be acknowledged. Cortical organoids resemble early stages of human brain development and therefore may not fully recapitulate the cellular composition and structural complexity of the mature brain exposed to radiotherapy. Hence, caution is required when translating these findings directly to radiation injury in adult patients. However, this developmental context may be particularly relevant for studying radiation effects in pediatric patients, where cranial irradiation during brain development is associated with progressive cognitive decline, along with IQ points lost per year. Standard cortical organoid culture does not develop a functional vascular network or a blood-brain barrier-like structure, hence vascular injury and drug pharmacological dynamics cannot be modeled directly. The absence of microglia limits the investigation of immune cell-driven synaptic remodeling and neuroinflammatory responses, both of which are crucial contributors to radiation-associated cognitive decline. In addition, variability in organoid maturation and cellular composition across different iPSC lines may impact both injury severity and recovery dynamics, highlighting the need for careful standardization and replication. Incorporating microglial components, vascularized assembloids, and functional assays such as calcium imaging or multielectrode recordings would further improve the physiological relevance of the model..

Overall, human iPSC-derived cortical organoids provide a scalable and mechanistically useful system for investigating CNS radiation injury and testing candidate mitigation strategies in a human-relevant context. Expanding the model to include microglia, vascularized assembloids, and long-term functional assessments would further enhance its physiological relevance. Lastly, comparison of the molecular changes observed in organoids after radiation to clinical biomarker data may ultimately help guide therapeutic approaches aimed at preserving neural integrity and cognitive function following cranial irradiation.

## Abbreviation

CNS: Central nervous system
hiPSCs: Human induced pluripotent stem cells
NSPP: 1-[(4-Nitrophenyl)sulfonyl]-4-phenylpiperazine
ROCK: Rho-associated coiled-coil containing protein kinase
TGF-β: Transforming Growth Factor-β
Wnt: Wingless-related integration site
EBs: Embryoid bodies
GFAP: Glial Fibrillary Acidic Protein
Tuj1: Neuron-specific β-III tubulin
CTIP2: COUP-TF-interacting protein 2
SATB2: Special AT-rich sequence-binding protein 2
MAP2: Microtubule-Associated Protein 2
S100β: S100 calcium-binding protein β
Olig1: Oligodendrocyte transcription factor 1
Olig2: Oligodendrocyte transcription factor 2
PAX6: Paired box protein 6
SYN1: Synapsin 1
PSD1: Postsynaptic density protein 1
GAP43: Growth-Associated Protein 43
CCL2: C-C motif chemokine ligand 2, also known as Monocyte Chemoattractant Protein-1, MCP-1
TNF-α: Tumor Necrosis Factor alpha
IL-1β: Interleukin-1 beta
MMP2: Matrix Metalloproteinase 2
scRNA-seq: Single-cell RNA sequencing

## Author contributions

LH conceived the study and designed the experiments. LH performed the experiments and data analysis. FP analyzed the single-cell RNA sequencing data and provided supervision and guidance throughout the project. HIK and AB provided critical support and troubleshooting throughout the experimental work. LH wrote the original draft. All authors contributed to reviewing and editing the manuscript and approved the final version.

## Data and Material Availability

The sequencing data have been submitted to Gene Expression Omnibus and are available with the following Accession Number: GSE295097.

## Acknowledgements

We thank the UCLA BSCRC Human Pluripotent Stem Cell Core/Bank for providing the iPSC lines used in this study, the UCLA Advanced Light Microscopy/Spectroscopy Core for providing access to imaging equipment and technical support, and the UCLA BSCRC Sequencing Core for assistance with single-cell RNA sequencing services. We thank Daria Azizad and Antoni Martija from Dr. Bhaduri’s laboratory for their guidance and assistance with organoid culture and experiments.

