## supplementary materials for "Temporal Mapping of Radiation-Induced Neural Injury and Mitigation in Human Cortical Organoids"

**Table of contents:**


1. Supplementary Figure

2. Supplementary Tables

3. Supplementary Materials and Methods

### Supplementary Figure 1.

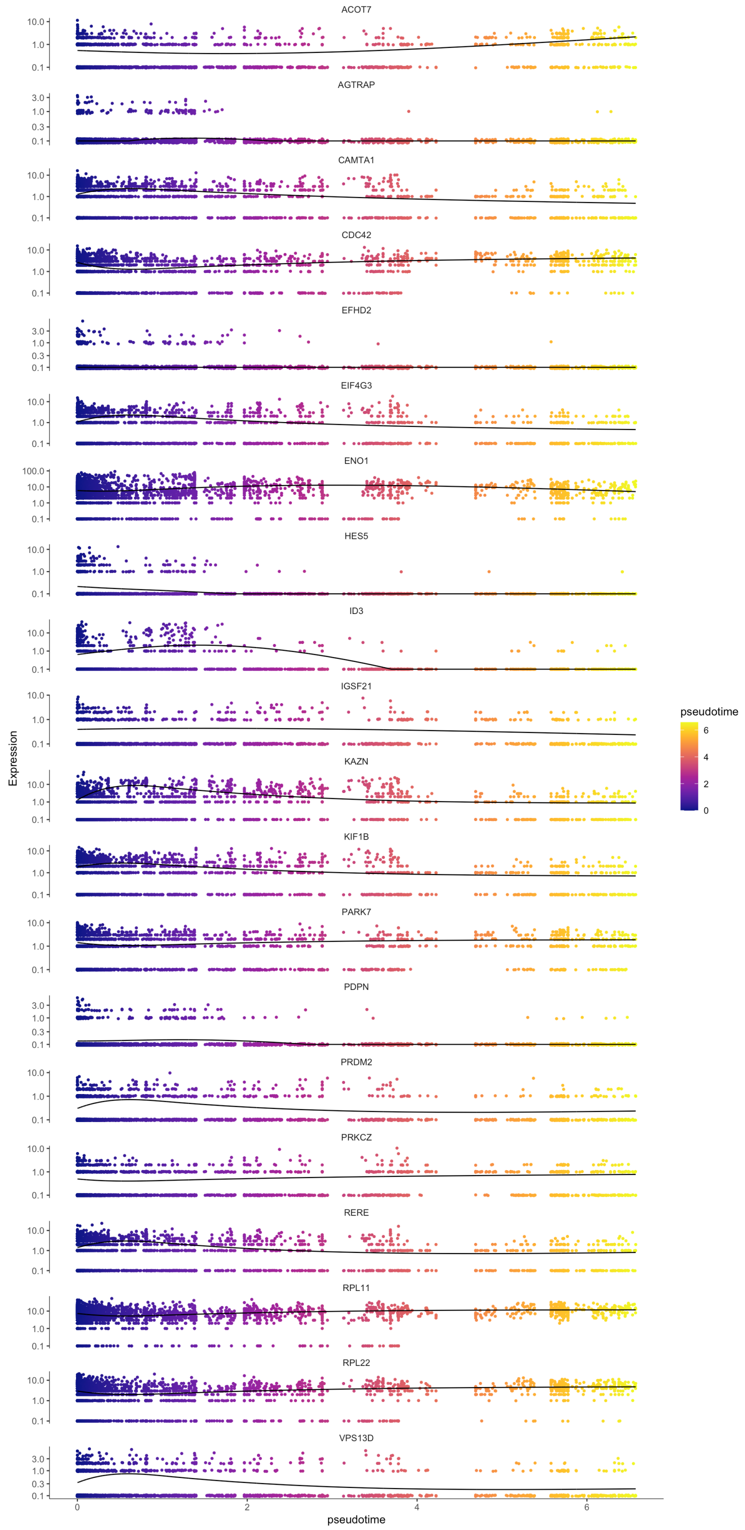


**Suppl. Figure 1.** Expression of Top 20 DEGs plotted along pseudotime inferred from the scVelo dynamic model.

**Supplemental Table 1. Human iPSC lines used in this study and their corresponding NIH hPSC registry information**

| Vendor | NIH Registry & Vendor Number |
|---|---|
| UCLA BSCRC | hiPSC2 |
| UCLA BSCRC | hiPSC19 |
| UCLA BSCRC | hiPSC21 |
| UCLA BSCRC | hiPSC22 |
| UCLA BSCRC | hiPSC23 |
| UCLA BSCRC | hiPSC27 |
| UCLA BSCRC | hiPSC28 |
| UCLA BSCRC | BSCRC23F1i1 |
| UCLA BSCRC | BSCRC23F1i3 |
| UCLA BSCRC | BSCRC39F2i11S |

BSCRC: Broad Stem Cell Research Center.

**Supplemental Table 2. PCR primer sequences (Integrated DNA Technologies) used for RT-PCR experiments.**

| Species | Gene name | Primer sequence (5'–3') |
|---|---|---|
| Human | NESTIN | Forward: GGTCTCTTTTCTCTTCCGTCC |
| | | Reverse: CTCCCACATCTGAAACGACTC |
| Human | SOX2 | Forward: CACACTGCCCCTCTCAC |
| | | Reverse: TCCATGCTGTTTCTTACTCTCC |
| Human | PAX6 | Forward: GCCCTCACAAACACCTACAG |
| | | Reverse: TCATAACTCCGCCCATTCAC |
| Human | TUBB3 | Forward: TTTGGACATCTCTTCAGGCC |
| | | Reverse: TTTCACACTCCTTCCGCAC |
| Human | MAP2 | Forward: CTTCAGCTTGTCTCTAACCGAG |
| | | Reverse: CTGCAACTATTCAAGGAAGTGG |
| Human | NEUN | Forward: GTAGAGGGACGGAAAATTGAGG |
| | | Reverse: CATAGAATTCAGGCCCGTAGAC |
| Human | GFAP | Forward: AGGACCTGCTCAATGTCAAG |
| | | Reverse: AGGCTGGTTTCTCGAATCTG |
| Human | S100β | Forward: CAGCAAGGAGACCAGGAAG |
| | | Reverse: TGGAAAACGTCGATGAGGG |
| Human | VIM | Forward: CGTGAATACCAAGACCTGCTC |
| | | Reverse: GGAAAAGTTTGGAAGAGGCAG |
| Human | OLIG1 | Forward: AGGCGCAATTCTCCAAGTG |
| | | Reverse: TGGACTGGAAGGTTCGAAATG |

| | | |
|---|---|---|
| Human | OLIG2 | Forward: AGCTCCTCAAATCGCATCC<br>Reverse: AAAAGGTCATCGGGCTCTG |
| Human | MBP | Forward: CCCAAGATGAAAACCCCGTAG<br>Reverse: TCCTCTCCCCTTTCCCTG |
| Human | TBR1 | Forward: GGAGCTTCAAATAACAATGGGC<br>Reverse: GAGTCTCAGGGAAAGTGAACG |
| Human | CTIP2 | Forward: ATGCCAGAATAGATGCCGG<br>Reverse: CTCTATCTCCAGACCCTCGTC |
| Human | SATB2 | Forward: AGGAGATGAAAAGGGCCAAG<br>Reverse: ACGGATGGTACAGAGGTTTTC |
| Human | SYN1 | Forward: CCCCAATCACAAAGAAATGCTC<br>Reverse: ATGTCCTGGAAGTCATGCTG |
| Human | PSD95 | Forward: CCGCTACCAAGATGAAGACAC<br>Reverse: TCCTCGTATTCCATCTCCCC |
| Human | GAP43 | Forward: TGACCAAGAACATGCCTGAAC<br>Reverse: GAAAGGACAGACTCACAGACG |
| Human | CCL2 | Forward: CAGAAGTGGGTTCAGGATTCC<br>Reverse: ATTCTTGGGTTGTGGAGTGAG |
| Human | MMP2 | Forward: ACCCATTTACACCTACACCAAG<br>Reverse: TGTTTGCAGATCTCAGGAGTG |
| Human | TNFα | Forward: ACTTTGGAGTGATCGGCC<br>Reverse: GCTTGAGGGTTTGCTACAAC |
| Human | IL-1β | Forward: ATGCACCTGTACGATCACTG |

| | | |
|---|---|---|
| | | Reverse: ACAAAGGACATGGAGAACACC |
| Human | PPIA | Forward: ATGCTGGACCCAACACAAAT<br>Reverse: TCTTTCACTTTGCCAAACACC |

**Supplemental Table 3. Percentage Composition for single-cell RNA sequencing.**

| Condition | Control | RT_72h | RT_2wk |
|---|---|---|---|
| Astrocyte-like | 0.017193948 | 0.013494318 | 0.026004728 |
| Dividing | 0.045392022 | 0.002840909 | 0.004728132 |
| Endothelial-like | 0.008940853 | 0.090198864 | 0.026004728 |
| Excitatory Neuron-like | 0.526134801 | 0.660511364 | 0.65248227 |
| Inhibitory Neuron-like | 0.03851444 | 0.02059659 | 0.03782506 |
| IPC-like | 0.02682256 | 0.00213068 | 0.00472813 |
| Newborn neuron-like | 0.082530949 | 0.002840909 | 0.007092199 |
| OPC-like | 0.034387895 | 0.010653409 | 0.016548463 |
| RG-like | 0.189821183 | 0.196022727 | 0.208037825 |
| RG.div-like | 0.030261348 | 0.000710227 | 0.016548463 |

RT: fractionated radiation (5 x 2 Gy); IPC: intermediate progenitor cell; OPC: oligodendrocyte progenitor cell; RG: radial glial cells; RG.div: dividing radial glial cells.

### Supplementary Materials and Methods

*Stem Cell Culture and Maintenance*

Human iPSCs were cultured on Matrigel-coated 6-well plates using mTeSR Plus medium supplemented with mTeSR Supplement (#100-0276, Stem Cell Technologies, Vancouver, Canada) and 1% Penicillin-Streptomycin. Media were refreshed every other day, and cells were passaged once they reached >80% confluency. For passaging, ReLeSR (#100-0483, Stem Cell Technologies) was applied to wells for 1 minute at room temperature and aspirated, followed by a 5-minute incubation at 37°C to promote detachment. Cells were gently dissociated into small clusters, avoiding complete single-cell dissociation, and transferred to new Matrigel-coated wells at a split ratio of 1:4 or 1:6. For cryopreservation, the passaging procedure was followed up to the dissociation step, after which cells were resuspended in 1 mL of CryoStor CS10 (#100-1061, Stem Cell Technologies) per well. The cell suspension was transferred into cryovials, initially stored at -80°C for 24-48 hours, and subsequently moved to liquid nitrogen for long-term storage. Cells were routinely screened for mycoplasma contamination using the Mycoplasma Detection Kit (#G238, Applied Biological Materials, Ferndale, WA).

*Cortical Organoid Generation*

Cortical organoids were generated using a modified protocol based on the previous publication [27], with adaptations from more recent studies [24, 26, 28]. Briefly, human iPSCs cultured to over 80% confluency in 6-well plates were dissociated by adding 1 mL of Accutase (#25-058-CI, Fisher Scientific, Waltham, MA) per well and incubated at 37°C for 5 minutes. To each well, 1 mL of Media 1 (M1) was added, consisting of GMEM

(#11710-035, LifeTech, Carlsbad, CA), 20% KnockOut Serum Replacement (#10828-028, LifeTech), 0.1 mM β-mercaptoethanol (#21985023, Sigma, St. Louis, MO), 1X Non-Essential Amino Acids (NEAA), 1X Sodium Pyruvate, and 1X Penicillin/Streptomycin (Fisher Scientific). Cells were gently scraped, transferred to 15 mL tubes, and centrifuged at 300 × *g* for 5 minutes. Pellets were resuspended in M1 containing three small molecules: Y27632 (20 µM, ROCK inhibitor), SB431542 (5 µM, TGF-β inhibitor), and IWR-1-endo (3 µM, Wnt inhibitor). Approximately 1 million cells were plated in 10 mL of M1 with small molecules into a 96-well V-bottom low-attachment plate (#277143, LifeTech) and allowed to aggregate for 72 hours without disturbance. On day 3, a partial medium change was performed, 50 µL was removed and replaced with 100 µL of fresh M1 containing the same small molecules to preserve organoid integrity. Full medium changes continued every other day. From day 7 onward, ROCK inhibitor was omitted from the formulation. On day 18, organoids were transferred to low-attachment 6-well plates (Fisher Scientific 07-200-601) and cultured with Media 2 (M2), composed of DMEM/F-12 with Glutamax (#10565-018, LifeTech), 1X N-2 supplement (#17502-048, LifeTech), 1X Lipid Concentrate (#11905-031, LifeTech), and 1X Penicillin/Streptomycin. M2 was used through day 35 with medium changes every other day. From day 35, Media 3 (M3) was introduced, consisting of DMEM/F-12 with Glutamax, 1X N-2 supplement, 1X Lipid Concentrate, 1X Penicillin/Streptomycin, 10% Fetal Bovine Serum (GIBCO, Grand Island, NY), 5 µg/mL Heparin (#H3149, Sigma), and 0.5% growth factor-reduced Matrigel. Starting on day 70, Media 4 (M4) was used, which included all M3 components with the addition of 1X B-27 Supplement with Vitamin A and insulin (LifeTech), and increased Matrigel concentration to 1%.

*Western blotting*

Equal amounts of protein were loaded onto 10% SDS-PAGE gels and subjected to electrophoresis for 2 hours. Samples were then transferred onto 0.45 μm nitrocellulose membrane (Bio-Rad, Hercules, CA) and blocked in 1x TBST containing 5% bovine serum albumin (BSA) for 30 minutes at RT, followed by incubation with primary antibodies against Neurofilament-L (#2837S, 1:1000, Cell Signaling Technology), GFAP (#12389S, 1:1000, Cell Signaling Technology), βIII-tubulin (#5568S, 1:1000, Cell Signaling Technology), and β-actin (#3700S, 1:1000, Cell Signaling Technology) in 1X TBST containing 5% BSA overnight at 4°C with gentle rocking. Membranes were then washed three times for 5 minutes each with 1X TBST and incubated with secondary antibodies, 1:5000 anti-mouse or anti-rabbit horseradish peroxidase (HRP; Cell Signaling Tchnology) in 1X TBST for two hours at RT with gentle rocking. Membranes were washed again three times for 5 minutes each with 1X TBST. Pierce ECL Plus Western Blotting Substrate (Thermo Fisher) was added to each membrane and incubated at RT for 5 minutes. The blots were then scanned using the Odyssey Fc imaging system (LI-COR Biosciences, Lincoln, NE). β-actin was used as a loading control. Densitometry was performed using ImageJ. The ratio of the protein of interest over its endogenous control was calculated and expressed as relative intensity.

*Immunofluorescence staining*

Human cortical organoids were fixed in 4% paraformaldehyde for 45 minutes at room temperature, followed by thorough rinsing with PBS. The fixed organoids were then equilibrated in 30% sucrose in PBS overnight at 4°C. After equilibration, organoids were embedded in a 1:1 mixture of OCT compound (Tissue-Tek, VWR, Radnor, PA) and 30% sucrose using an OCT embedding mold (#14-373-65, Fisher Scientific) and rapidly frozen on dry ice. Embedded organoids were either stored at -80°C for up to 3 months or immediately cryosectioned at a thickness of 10-16 μm onto glass slides for subsequent staining.

For immunofluorescence staining, sections were rinsed in PBS for 15 minutes and subjected to antigen retrieval using a citrate-based unmasking solution (10 mM sodium citrate, pH 6, Vector Labs, Newark, CA), heated to 95°C for 20 minutes. Following antigen retrieval, sections were permeabilized and blocked for 30 minutes at room temperature using a blocking buffer composed of 5% goat serum, 1% bovine serum albumin (BSA), and 0.1% Triton X-100 in PBS. Primary antibody incubations were performed overnight at 4°C in blocking buffer, using the following antibodies and dilutions: cleaved caspase-3 (rabbit, 1:400, Cell Signaling technology, #9661t, Danvers, MA), Nestin (mouse, 1:1000, Cell Signaling Technology, #33475s), Pax6 (rabbit, 1:400, Cell Signaling Technology, #60433t), βIII-tubulin (mouse, 1:1000, Abcam, ab78078), GFAP (rabbit, 1:400, Cell Signaling Technology, #80788s), Ki67 (rabbit, 1:500, Cell Signaling Technology, #11882s), MBP (rabbit, 1:400, Cell Signaling Technology, #78896t), γH2AX (rabbit, 1:500, Cell Signaling Technology, #9718s), CTIP2 (rabbit, 1:400, Cell Signaling Technology, #12120t), SATB2 (mouse, 1:500, Abcam, ab51502). The following day, sections were incubated with species-specific secondary antibodies: Alexa Fluor 594-conjugated goat anti-rabbit immunoglobulin G (IgG) (H/L) antibody (1:1,000 (Invitrogen)) and Alexa Fluor 488-conjugated goat anti-mouse IgG (H/L) antibody (1:1,000

(Invitrogen)), followed by nuclear counterstaining with Hoechst33342 (Thermo Fisher). Slides were mounted using ProLong™ Gold antifade reagent (Invitrogen) and stored at 4°C until imaging. Fluorescent images were acquired using a confocal microscope (Nikon A1, Melville, NY).
